# Online milk composition analysis with an on-farm near-infrared sensor

**DOI:** 10.1101/2020.06.02.129742

**Authors:** Jose A. Diaz-Olivares, Ines Adriaens, Els Stevens, Wouter Saeys, Ben Aernouts

## Abstract

On-farm monitoring of milk composition can support close control of the udder and metabolic health of individual dairy cows. In previous studies, near-infrared (NIR) spectroscopy applied to milk analysis has proven useful for predicting the main components of raw milk (fat, protein, and lactose). In this contribution, we present and evaluate a precise tool for online milk composition analysis on the farm. For each milking, the online analyzer automatically collects and analyses a representative milk sample. The system acquires the NIR transmission spectra of the milk samples in the wavelength range from 960 to 1690 nm and performs a milk composition prediction afterward.

Over a testing period of 8 weeks, the sensor collected 1165 NIR transmittance spectra of raw milk samples originating from 36 cows for which reference chemical analyses were performed for fat, protein, and lactose. For the same online sensor system, two calibration scenarios were evaluated: training post-hoc prediction models based on a representative set of calibration samples (*n* = 319) acquired over the entire testing period, and training real-time prediction models exclusively on the samples acquired in the first week of the testing period (*n* = 308).

The obtained prediction models were thoroughly tested on all the remaining samples not included in the calibration sets (*n* respectively 846 and 857). For the post-hoc prediction models, this resulted in an overall prediction error (root-mean-squared error of prediction, RMSEP) smaller than 0.08% (all % are in *w*/*w*) for milk fat (range 1.5-6.3%), protein (2.6-4.3%) and lactose (4-5.1%), while for the real-time prediction models the RMSEP was smaller than 0.09% for milk fat and lactose, and smaller than 0.11% for protein. The milk lactose predictions could be further improved by taking into account a cow-specific bias. The presented online sensor system using the real-time prediction approach can thus be used for detailed and autonomous on-farm monitoring of milk composition after each individual milking, as its accuracy is well within the ICAR requirements for on-farm milk analyzers and even meet the ICAR standards for laboratory analysis systems for fat and lactose. For this real-time prediction approach, a drift was observed in the predictions, especially for protein. Therefore, further research on the development of online calibration maintenance techniques is required to correct for this model drift and further improve the performance of this sensor system.

## Introduction

The metabolism of dairy cows is heavily conditioned by milk production. As a result, milk composition can inform about the cow’s nutritional, metabolic, and health status (McParland et al., 2014). In standard dairy practices, cows are milked twice or three times a day, which implies that milk samples can be taken and analyzed regularly without interfering in the animal’s daily life. Therefore, frequent analysis of the produced milk can be considered a very efficient way to monitor the performance, efficiency, and welfare of individual dairy cows.

Critical dairy health issues such as ketosis and mastitis produce changes in the concentration of the main components of raw milk (fat, protein, and lactose). For example, mastitis causes a decrease in lactose concentration (Bobbo et al., 2016; Forsbäck et al., 2010, 2009; Gonçalves et al., 2016), while ketosis is associated with increased fat and decreased protein contents (Brandt et al., 2010). Frequent and accurate monitoring of these changes over time allows for detecting alterations of the udder and metabolic health of individual cows (Mäntysaari et al., 2019).

Visible (Vis) and near-infrared (NIR) spectroscopy is a promising technique for the on-farm monitoring of the main components of raw milk, a potential scrutinized by numerous researchers in the past (Kawamura et al., 2007; Kawasaki et al., 2008; Melfsen et al., 2012; Tsenkova et al., 1999). Several commercially available technologies make use of this technique, mainly in the Vis and short-wave NIR wavelength range (400 to 1000 nm). Despite the low cost of detectors for that wavelength range, the performance in this region often provides unsatisfactory results for the prediction of milk composition (Fadul-Pacheco et al., 2018; Kaniyamattam and De Vries, 2014). This suboptimal performance is apparent when compared to NIR analyses performed using the long-wave NIR (LW-NIR) range from 1000 to 1700 nm (Aernouts et al., 2011).

Both reflectance and transmittance spectra can be collected using LW-NIR spectroscopy. In previous works, LW-NIR transmittance has shown an exceptional performance for raw milk composition prediction, with a root-mean-square error (RMSE) lower than 0.13% (*w*/*w*) and a coefficient of determination (*R*^2^) higher than 0.9 for fat, protein and lactose predictions (Aernouts et al., 2011; Saranwong and Kawano, 2008; Tsenkova et al., 1999).

Using LW-NIR reflectance, fat and protein show a strong prediction performance, as fat globules and casein micelles are strongly linked to the NIR scattering. These scattering phenomena are caused by their physical structures, in addition to the absorbance of NIR radiation by their covalent bonds (Bogomolov and Melenteva, 2013). In contrast, lactose predictions are relatively poor (RMSE higher than 0.15% and an R^2^ lower than 0.75), as the majority of the reflected photons acquired by the sensor have limited interaction with the milk serum, in which lactose is diluted (Aernouts et al., 2015b).

Despite the potential of LW-NIR spectroscopy shown in previous studies, the strong absorption of LW-NIR radiation by water molecules and the severe light scattering by fat globules and casein micelles pose significant challenges for its on-farm use. For example, the thickness of the sample can only be 1 to 2 mm in order to obtain sufficient LW-NIR transmittance (Aernouts et al., 2011; Tsenkova et al., 1999), which complicates its in-line implementation due to its impact on the flow of the milk acquisition process on milking systems. This implies that a representative milk sample needs to be taken using a bypass, as demonstrated by Kawasaki et al. (Kawasaki et al., 2008).

Although many researchers have already shown the potential of LW-NIR spectroscopy for milk composition analysis in the laboratory, studies testing this technology under farm conditions are scarce and often include only a limited number of milk samples measured over a few test days (Kawasaki et al., 2008; Melfsen et al., 2012). Moreover, all previous studies, even the ones performed on farms, have made use of a post-hoc approach to predict the milk composition, in which the samples to train and test the prediction models were posteriorly selected from the complete dataset, both covering the full variability and time span of that dataset. None of these previous studies have considered a real-time prediction of the milk composition using calibration models that were already established before an independent set of validation samples was measured.

In this study, we present an on-farm sensor system making use of LW-NIR spectroscopy to monitor the milk composition for each individual cow and every milking session. The goal of this work is to evaluate the accuracy of this tool and its potential for online measurement of the milk fat, protein, and lactose concentrations, with a real-time prediction approach. Moreover, to investigate model drift, the real-time predictions are compared to the ones obtained with the post-hoc approach followed in former studies. Additionally, it was investigated whether a cow-specific bias correction can further improve the performance of the real-time prediction models.

## Materials and methods

### Sensor system to measure the long-wave near-infrared transmittance of raw milk

The sensor system, illustrated in Figure 1, consists of (a) a NIR spectrometer (1.7-256 Plane Grating Spectrometer, Carl Zeiss, Jeda, Germany) with a cooled InGaAs (indium-gallium-arsenide) diode array of 256 pixels (2.86 nm pixel resolution in the 960 to 1690 nm range); (b) a 20 Watt halogen light source (998079-14, Welch Allyn, New York, USA); (c) an optical measuring unit and (d) a dedicated computer which runs the control and spectral acquisition software in LabView 2012 (National Instruments, Austin, USA). All previous elements are contained in a compact and temperature-controlled (PowerCool, Laird, Liberec, Czech Republic) steel housing, which protects them from ammonia, dust, and moisture.

**Figure 1:**
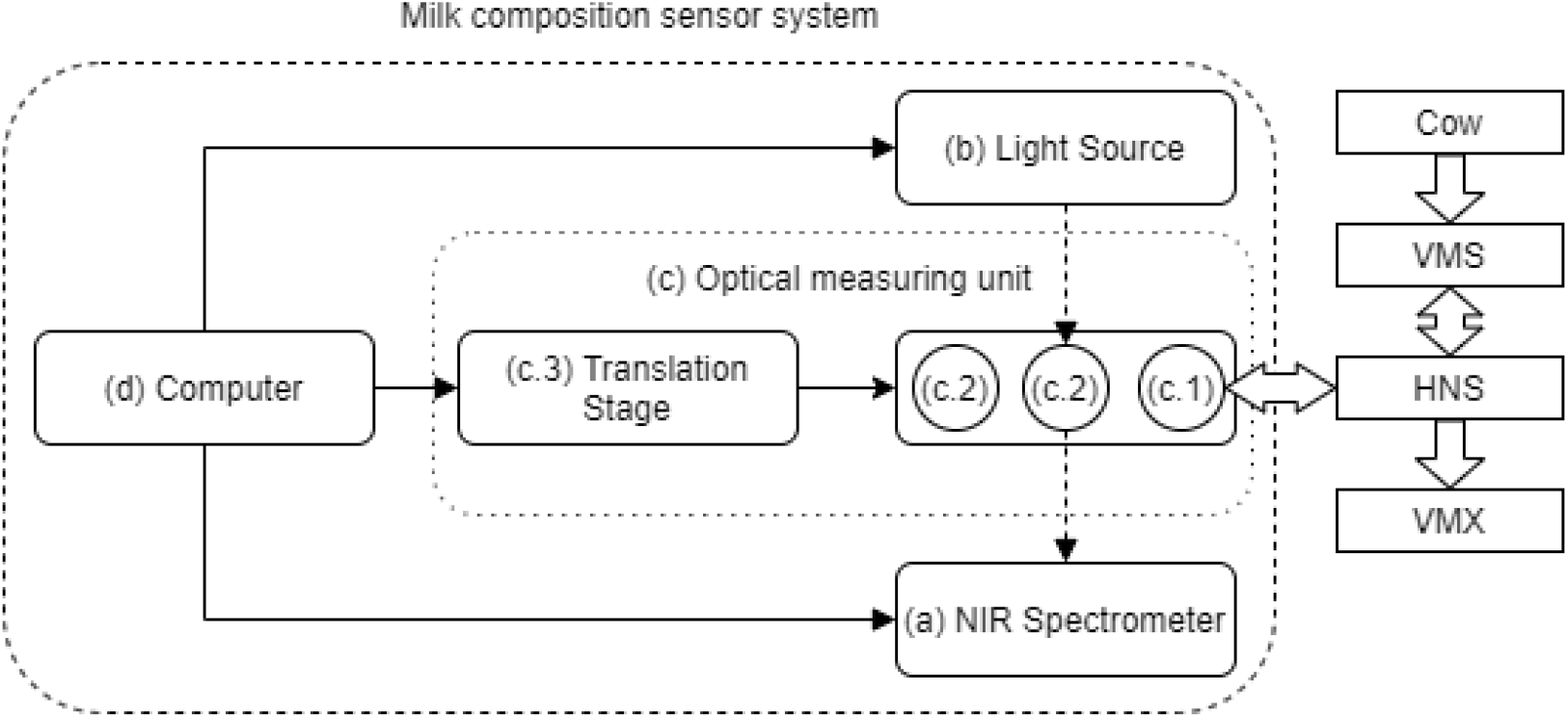
Conceptual depiction of the sensor system. Bold black arrows indicate control flow, while dashed black arrows indicate light flow from the source. White block arrows illustrate milk flow.

The optical measuring unit comprises (c.1) a round borosilicate cuvette with an inner thickness and diameter of respectively 2 and 26 mm and containing 1.2 mL of raw milk during spectral sample measurements; (c.2) a spectral reference pair with a 2 mm thick Spectralon (Labsphere, North Sutton, USA) white standard and a 1% Acktar Black (Acktar, Hohenaspe, Germany) coated plate as a dark standard, and (c.3) a translation stage, consisting of a filter slide (FS40, OWIS, Frankfurt, Germany) which was automated with a stepper motor (L4118L1804, Nanotec, Munich, Germany). This translation stage holds the cuvette and the spectral reference pair and allows to alternate between them in order to perform spectral reference measurements before measuring the milk sample. The transmitted light is collected by a converging-type lens with a focal length of 31.6 mm (770303-9020-000 2:1, Carl Zeiss, Jena, Germany) and guided to the spectrometer by an optical fiber (600 μm core diameter, low-OH). The lens is aligned with the center of the halogen light source and the translation stage moves perpendicularly to the lens-halogen axis when alternating between the cuvette and the white and dark spectral references. When the dark spectral reference is in position, the stray light and background noise of the spectrometer are quantified.

For each milk sample in the cuvette, the NIR spectrometer recorded LW-NIR spectra in transmittance mode in the wavelength range from 960 to 1690 nm. To obtain a high signal-to-noise ratio, these spectra were acquired with an integration time of 100 ms collecting 100 repeated measurements per milk sample, aggregated into an average spectrum for each sample. In addition, dark and white reference spectra were acquired using the same settings while the milk sample was loaded in the cuvette, just before performing a spectral recording for that milk sample.

### Experimental setup and standardized chemical analysis of milk samples

The sensor system was tested in combination with a VMS™ Classic (DeLaval, Tumba, Sweden) automated milking system (AMS) at the experimental dairy farm ‘Hooibeekhoeve’ of the province of Antwerp (Geel, Belgium). A Herd Navigator™ sampler (HNS Supra+, DeLaval) was installed on the AMS collecting representative milk samples of about 300 mL of each milking. With this sampler, a fraction (±100 mL) of the collected milk samples can be pumped to the Herd Navigator™ analyzer (DeLaval) for milk biomarker analysis, while another fraction (±30 mL) can be sent to the VMX™ (DeLaval) to take a milk sample for laboratory milk analysis in the context of milk recording programs. The sensor system was installed as a bypass on the backflow of the Herd Navigator™ sampler and received at least 150 mL of raw milk. Thanks to thorough sample mixing in the Herd Navigator™ sampler, the composition of the milk analyzed by the sensor system and the milk taken by the VMX™ milk sampler can be assumed identical.

Over a period of 8 weeks, the sensor system analyzed raw milk originating from 41 Holstein cows (lactation stage 168 ± 84 DIM, parity 2.0 ± 1.1) on an irregular basis, dependent on when a milk sample was taken for reference chemical analysis in the context of an unrelated feeding trial. The periods for which milk samples were collected varied between 21 and 85 consecutive hours per week (i.e., between 59 and 316 samples, with an average of 159 samples per week). The milk samples taken by the VMX™ were preserved with bronopol (0.3 mg/mL) and analyzed at the Milk Control Center (MCC Vlaanderen, Lier, Belgium) within three days after sample collection. The reference chemical analysis (fat, protein, and lactose) of each sample was performed according to ISO 9622 (ISO, 2013) with a Milkoscan FT+ (Foss A/S, Hillerød, Denmark).

In total, milk samples and spectral measurements corresponding to 1270 individual milking sessions were collected during the experimental period. Fifty-nine of them originated from four dairy cows that calved during the trial and from one individual animal that was less than five days in milk at the beginning of the experiment. To avoid the inclusion of colostrum samples, they were removed from the dataset (Abd El-Fattah et al., 2012). Forty-six additional samples had to be discarded as a consequence of unsuccessful reference chemical analyses. The final dataset consisted of 1165 raw milk samples from 36 cows for which both transmission spectra and reference chemical analyses were available.

### Selection and normalization of the sample spectra

The dataset with transmittance spectra and the results from the reference chemical analyses were imported in R version 3.5.1 (R Core Team, 2018) to be analyzed with a custom chemometrics toolbox (Aernouts et al., 2020).

By default, a pair of dark and white reference spectra was acquired every time a milk sample spectrum was recorded. This pair of reference spectra was used to normalize the sample spectra according to the formula 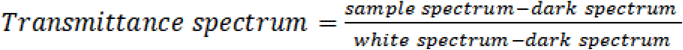. This procedure corrects the spectrum of each sample for drift in the spectral sensitivity of the spectrometer and the intensity spectrum of the light source, both mainly influenced by stray light or environmental temperature changes (Andersen et al., 2013).

This correction is only needed when the drift effect is higher than the stochastic noise introduced by the acquisition of the reference spectra (Slutsky, 1997). Therefore, to identify degradation in the prediction caused by this drift and accordingly improve the milk composition predictions, different time intervals for acquiring a new pair of dark and white reference spectra were evaluated. The procedures to obtain an optimal reference acquisition interval are detailed in the supplementary materials of this publication.

Based on this analysis, a new spectral reference pair was considered right after connecting the sensor system to the AMS and every 0.5 hours after this initialization.

### Development and validation of the post-hoc prediction models

In previous studies, a post-hoc prediction approach was followed in which the samples to train and test the prediction models were selected from a single dataset, both covering the full variability and time span of that dataset. For an objective comparison, a similar approach was initially followed in this study too. To this end, the duplex algorithm (Snee, 1977) was used to divide the total dataset of the milk transmittance spectra and their respective reference chemical analyses into an elaborate and representative calibration set of approximately 300 samples (Shetty et al., 2011) and a test set containing the remainder of the samples. Not taking into account the time when the samples were measured, the post-hoc approach uses the Mahalanobis distance between the samples in the 3-dimensional space of the reference chemical analyses (fat, protein, and lactose) to obtain two subsets with similar descriptive statistics and correlations for these milk components. To prevent over-optimistic test results due to modeled cow-specific effects, all samples from the same cow were grouped either in the calibration or test set (Kemps et al., 2010).

The wavelength range in the spectra where the absorption by water molecules is predominant (1360 to 1500 nm), and thus the transmittance signals are very low, were removed to reduce the introduction of spectral noise. The transmittance spectra of the calibration samples were used to train their respective partial least squares regression (PLSR) models to predict the individual milk components. The optimal number of latent variables for each PLSR model, the best spectral preprocessing, and the most informative spectral wavelengths were selected based on the model performances in group-wise cross-validation for the samples in the calibration set (Aernouts et al., 2020). To this end, the calibration set was split into ten groups, each containing all the spectra of a single cow. Accordingly, all samples from the same cow were either included or excluded from the training set to prevent modeling cow-specific effects or relations (Kemps et al., 2010). The least complex model (with the lowest number of latent variables) that did not perform significantly worse than the model with the lowest root-mean-squared error of cross-validation (RMSECV) was selected (Haaland and Thomas, 1988). This comparison was based on one-sided paired t-tests (*α* = 0.05) applied to the absolute residuals of the cross-validated calibration samples (Cederkvist et al., 2005). This procedure was repeated for the three different milk components, obtaining a prediction model per component. As the resulting models were trained on samples extracted from the entire experimental period of 8 weeks, they are hereafter referred to as the post-hoc prediction models. Using these models, milk composition predictions were obtained for the samples in the test set, and the respective residuals were derived. These errors were then combined into a root-mean-square error of prediction (RMSEP).

### Development and validation of the real-time prediction models

To evaluate the performance of the sensor system in predicting milk composition after each individual milking, which is the main goal of this study, prediction models for fat, protein, and lactose were trained on samples taken in the first week only (*n* = 308) and validated on all subsequent samples, which were measured over the following seven weeks (*n* = 857). These models are hereafter referred to as the real-time prediction models. The procedures to train, test, and compare the models were identical to the ones previously described for the post-hoc prediction models.

To compare the prediction performance of the real-time and the post-hoc approaches, they were compared using a statistical test on the squared residuals of the milk component predictions for the samples in the test set. As the test sets of the real-time and the post-hoc models did not entirely overlap, an unpaired one-way analysis of variance (ANOVA), with the model type as a two-level (post-hoc or real-time) factor, was performed. The two approaches were compared mutually with an HSD multiple comparisons test (*α* = 0.05) when the ANOVA procedure (*α* = 0.05) indicated a significant effect of the model type.

### Analysis of the cow-specific bias on the milk component predictions by the real-time models

To evaluate whether bias was present in the milk composition predictions for specific dairy cows, an analysis of this phenomenon was undertaken (Anderson et al., 2016). Over time, this bias can be observed as a cow-specific baseline in the residuals that are obtained after subtracting the predicted milk compositions from the respective reference chemical analyses. For the real-time models, all cows were represented in the calibration set, as well as the test set. This was not true for the post-hoc models, due to how the duplex procedure was implemented to split the original dataset. Accordingly, the cow-specific bias on the three milk component predictions was only studied for the real-time models.

The median value of the residuals of the samples in the calibration set was calculated for each individual cow. First, the agreement between those medians and the prediction residuals of the samples in the test set was studied for all cows using the concordance correlation coefficient (Lin, 1989). Additionally, the cow-specific bias obtained from the calibration set was then subtracted from all further milk composition predictions for that same animal in the test set. Next, the performance of the real-time model for each component was evaluated before and after the correction of the cow-specific bias. This comparison was made with a two-way repeated-measures ANOVA analysis applied on the squared residuals of the samples in the test set, with “cow-specific bias correction” as a fixed factor with two levels (affirmative or negative) and the sample number as a random factor. Only when a significant effect of the correction was detected by the ANOVA procedure (*α* = 0.05), an HSD multiple comparisons test (*α* = 0.05) was used to test whether the correction for the cow-specific bias resulted in an improvement.

## Results and discussion

### Collection of long-wave near-infrared spectra

In Figure 2, the absorbance spectra, obtained by the logarithmic transformation of the inverse of the transmittance spectra, is illustrated as solid blue lines for all the milk samples of a single dairy cow. It presents the variability in the observations acquired from the same animal across the duration of the experiment. Additionally, the maximum and minimum spectra of the entire dataset are shown as solid red lines. Clear absorbance peaks can be observed around 970, 1200 and 1450 nm. The absorbance units between 4 and 5 for the latter peak indicate that only a very small fraction (10^-4^ – 10^-5^) of the incident photons around this wavelength range was transmitted, resulting in a low signal to noise ratio.

**Figure 2.**
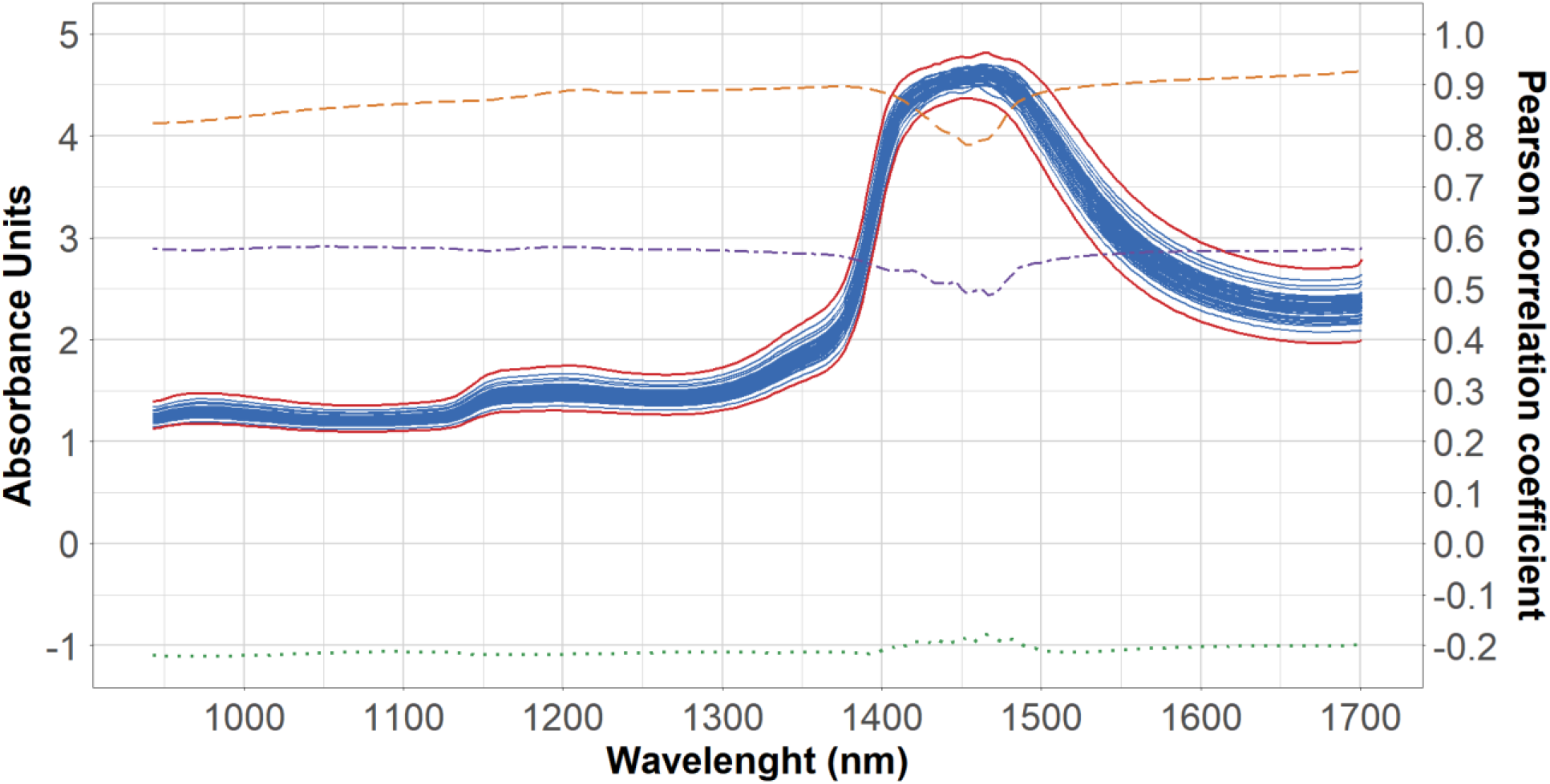
Absorbance spectra for a single dairy cow (solid blue lines) collected over a period of 8 weeks. Minimum and maximum (bold red solid lines) absorbance spectral values for the entire dataset are depicted. The Pearson correlation coefficients illustrate for every wavelength the linear correlation between the near-infrared absorbance by the 1165 raw milk samples and their fat (orange dashed line), protein (purple dot-dashed line), and lactose (green dotted line) concentration.

The high correlation between the fat content and the absorbance values over the full wavelength range can be attributed to the scattering of light by the fat globules. As a higher fat content results in stronger light scattering, it reduces the transmittance and thus increases the absorbance, resulting in a nearly constant correlation between the fat content and the NIR absorbance over the full wavelength range. In Figure 2, the maximum and minimum absorbance spectra (bold red solid lines) of the entire dataset corresponded to respectively the maximum (6.25%) and the minimum fat percentage (1.54%). An absorption band of C-H bond overtone, characteristic of fat molecules, can be observed as a slight increase in the correlation with the fat content of the NIR absorption around 1210 nm (Zamora-Rojas et al., 2013). A significant positive correlation between protein and the spectra can be observed. This relationship can be mainly attributed to the correlation between the protein and the fat content (*r* = 0.55), a relationship associated with their genetic correlation (Rosati and Van Vleck, 2002). In contrast, a very weak negative correlation was found between milk lactose content and the NIR absorption spectra. Protein and lactose do not display clear absorption peaks, as they overlap with the more dominant absorption of water and fat.

### Development and validation of the post-hoc prediction models

Following the duplex algorithm, the original dataset of 1165 raw milk samples was split between a calibration set (*n* = 319) and a test set (*n* = 846). The descriptive statistics of the fat, protein, and lactose content of the resulting sets and their correlations are summarized in Table 1. The comparable average and variability show that the data splits are representative for the whole dataset, which is critical for developing a robust prediction model (Saeys et al., 2008). Comparing the statistical parameters of this dataset to the data from milk recording programs in Flanders (Aernouts et al., 2011), showed that our samples also have comparable values and variability as the cow population of this region.

**Table 1.** Basic statistics and Pearson correlations for the main milk components in the calibration and test sets used to construct the post-hoc prediction models.

| Component | Calibration (n = 319) |  |  |  |  |  | Test (n = 846) |  |  |  |  |  |
| --- | --- | --- | --- | --- | --- | --- | --- | --- | --- | --- | --- | --- |
|  | Basic statistics (% w/w) |  |  |  | Pearson corr. |  | Basic statistics (% w/w) |  |  |  | Pearson corr. |  |
|  | Mean | SD | Min | Max | Fat | Protein | Mean | SD | Min | Max | Fat | Protein |
| Fat | 3.54 | 0.92 | 1.54 | 5.81 | 1 | - | 3.5 | 0.76 | 1.54 | 6.25 | 1 | - |
| Protein | 3.44 | 0.34 | 2.75 | 4.26 | 0.51 | 1 | 3.42 | 0.34 | 2.63 | 4.34 | 0.57 | 1 |
| Lactose | 4.65 | 0.19 | 3.98 | 5.1 | -0.11 | -0.23 | 4.71 | 0.14 | 3.99 | 5.03 | -0.19 | -0.11 |

The details for the different time intervals that were evaluated for acquiring a new pair of dark and white are presented in the supplementary material, in tables S1 and S2. Based on this analysis, it was decided to take a new spectral reference pair at the start of each session after connecting the sensor system to the AMS and every 0.5 hours after this initialization. Based on the obtained results, it is expected that if new spectral references are taken with a higher frequency than this optimum, unnecessary stochastic noise, typical for spectral measurement, would be introduced. In contrast, if new spectral references are taken with a lower frequency than this optimum, significant drift in the spectral output of the light source and spectral sensitivity of the spectrometers might not be fully captured and accounted for, also negatively affecting the results.

Figure 3 shows the prediction performance of the post-hoc models for the calibration samples in cross-validation (red circles) and the test samples (green crosses). Very good predictions are obtained for all three milk components, with RMSEP values smaller than 0.08% (*w*/*w*). For fat and protein, more than 98% and 94% of the variation in the reference chemical analyses for these components can be explained by their respective prediction model. The lower *R*^2^ values observed for lactose (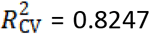 and 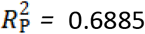) can be partly attributed to the limited range of this component in this particular dataset. Additionally, it can be observed (Figure 3c) that the majority of the test samples have a lactose concentration greater than 4.5% (*w*/*w*), while lactose calibration samples span a wider distribution, contributing to the discrepancy between the 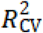 and 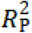. values while the RMSE values are very similar. This suggests that the split performed by the duplex algorithm had limited success for lactose. For both fat and lactose, RMSECV values are slightly higher than RMSEP, which can be partly attributed to the greater dispersion of the calibration data for both milk components (SD of 0.92% for fat and 0.19% for lactose) against the test data (SD of 0.76% for fat and 0.14% for lactose, all %*w*/*w*) and the higher presence of samples that are further away from the identity line in the calibration data compared to the test data. The PLSR model for lactose had the highest complexity (16 latent variables), which is considerably higher than the 3 and 10 latent variables for fat and protein, respectively.

**Figure 3.**
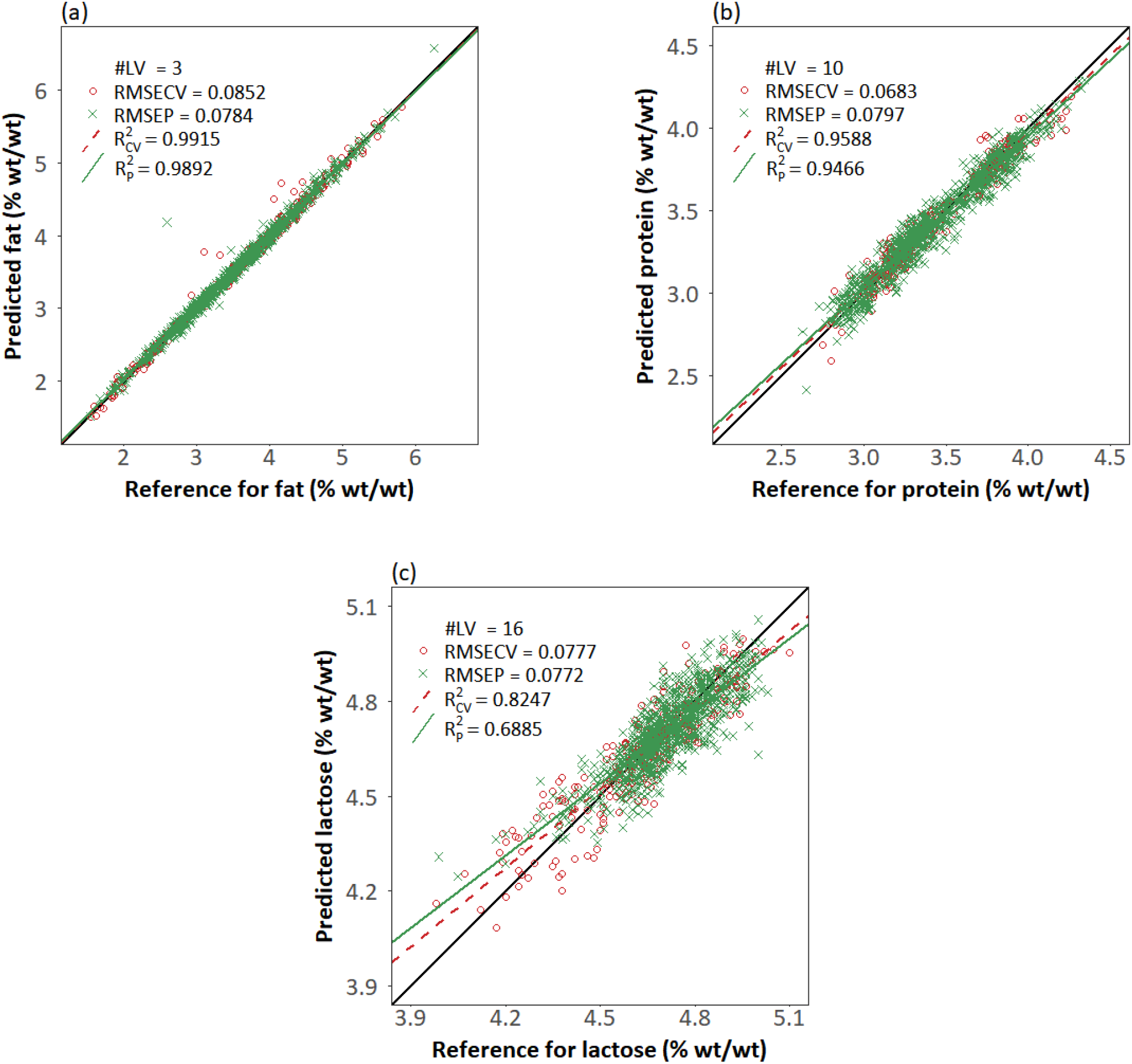
Scatter plots of the predicted versus reference milk composition content for the calibration set in cross-validation (red circles) and the test set (green crosses) for (a) fat, (b) protein and (c) lactose, for the post-hoc approach. #LV = number of latent variables used by the model; RMSECV = root-mean-square error of cross-validation; RMSEP = root-mean-square error of prediction; 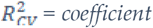 of determination for cross-validation; 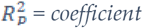 of determination for prediction. The black solid lines are the identity lines.

As can be seen in Table 2, the performance of these models is comparable to the results for the prediction of the raw milk composition based on LW-NIR spectra reported by other researchers. The selection of these works prioritized potential on-farm applications and the use of the LW-NIR wavelength range (Aernouts et al., 2011; Kawasaki et al., 2008; Melfsen et al., 2012; Saranwong and Kawano, 2008; Tsenkova et al., 2001). These studies were all performed in transmission mode, except for Melfsen et al., (2012), who used diffuse reflectance mode.

**Table 2.** Comparison of obtained model performance to the values reported by other researchers.

| Source | *n* | RMSEP (% w/w) | | | $R_P^2$ | | |
| --- | --- | --- | --- | --- | --- | --- | --- |
|  |  | Fat | Protein | Lactose | Fat | Protein | Lactose |
| Post-hoc model | 846 | 0.0784 | 0.0797 | 0.0772 | 0.9892 | 0.9466 | 0.6885 |
| Kawasaki et al., 2008 | 72 | 0.259<sup>Δ</sup> | 0.151<sup>Δ</sup> | 0.262<sup>Δ</sup> | 0.95 | 0.72 | 0.83 |
| Melfsen et al., 2012* | 262 | 0.09<sup>Δ</sup> | 0.05<sup>Δ</sup> | 0.061<sup>Δ</sup> | 0.998 | 0.98 | 0.92 |
| Tsenkova et al., 2001 | 86 | 0.261<sup>Δ</sup> | 0.132<sup>Δ</sup> | 0.084<sup>Δ</sup> | 0.98 | 0.45 | 0.72 |
| Aernouts et al., 2011 | 100 | 0.046 | 0.133 | 0.115 | 0.997 | 0.927 | 0.883 |
| Saranwong and Kawano, 2008<sup>γ</sup> | 47-43-46 | 0.03<sup>Δ</sup> | 0.072<sup>Δ</sup> | 0.096<sup>Δ</sup> | 0.99 | 0.96 | 0.89 |

RMSEP = Root-mean-square error of prediction; $R_P^2$ = coefficient of determination; *n* = number of samples in the test set.
<sup>Δ</sup>RMSEP values calculated from standard error of prediction (SEP), bias, and number of samples, provided by the authors, with the following relationship: $RMSEP = \sqrt{((n - 1)/n) \times SEP^2 + bias^2}$ .
\*In this study, diffuse reflectance was used, all other studies used transmittance.
<sup>γ</sup>Saranwong and Kawano (2008) used a different number of test samples for fat, protein, and lactose, respectively.

RMSEP and 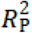 values were considered in the examination of the literature sources (Table 2). The analysis by Saranwong and Kawano (2008), Aernouts et al. (2011), and the post-hoc model achieved outstanding accuracy for fat prediction, with RMSEP values smaller than 0.08% (*w*/*w*). Saranwong and Kawano (2008), Melfsen et al. (2012) and the post-hoc approach of the present study are the best performing for predicting protein content, with RMSEP values that did not exceed 0.08% (*w*/*w*). Melfsen et al. (2012) and the post-hoc model in this study are very good for predicting lactose content in milk, with RMSEP values under 0.08 % (*w*/*w*).

In comparison to the results reported by other researchers, we obtained low RMSEP and high 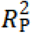 values, underlining the general suitability of the sensor system and calibration procedure for predicting the main constituents in raw milk.

However, by selecting calibration samples throughout the experiment, the literature sources and present post-hoc models do not provide an objective and complete image on how the system would perform in a real-time setting. In order to evaluate the capability of the on-farm sensor system to perform real-time predictions, the validation should be performed on samples that are measured after the initial calibration, thus being independent of the samples used for training the prediction models.

### Development and validation of the real-time prediction models

To test the potential for real-time prediction of the milk composition, the 308 raw milk samples collected in the first week of the trial were assigned to the calibration set, while those collected in weeks 2 to 8 (*n* = 857) were assigned to the test set. The descriptive statistics for the fat, protein, and lactose content of both sets and their correlations are summarized in Table 3. This shows that the samples of the first week are representative of the variation present in the samples of weeks 2 to 8.

**Table 3.** Descriptive statistics and Pearson correlations of the main milk components in the calibration and test set for the real-time prediction models.

| Component | Calibration (n = 308) |  |  |  |  |  | Test (n = 857) |  |  |  |  |  |
| --- | --- | --- | --- | --- | --- | --- | --- | --- | --- | --- | --- | --- |
|  | Basic statistics (% w/w) |  |  |  | Pearson corr. |  | Basic statistics (% w/w) |  |  |  | Pearson corr. |  |
|  | Mean | SD | Min | Max | Fat | Protein | Mean | SD | Min | Max | Fat | Protein |
| Fat | 3.44 | 0.86 | 1.61 | 6.25 | 1 | - | 3.54 | 0.79 | 1.54 | 5.72 | 1 | - |
| Protein | 3.45 | 0.35 | 2.65 | 4.24 | 0.53 | 1 | 3.42 | 0.34 | 2.63 | 4.34 | 0.57 | 1 |
| Lactose | 4.74 | 0.15 | 4.17 | 5.1 | -0.15 | -0.11 | 4.68 | 0.15 | 3.98 | 5.02 | -0.16 | -0.18 |

Identically to the post-hoc models, the selected time interval corresponded to taking a new spectral reference pair at the start of each session after connecting the sensor system to the AMS and every 0.5 hours after this initialization. These results are presented in the supplementary material (tables S3 and S4).

The prediction performances of the post-hoc and real-time approaches were mutually compared based on the residuals for the samples in the test set (Table 4). Since the test sets for those two approaches were not identical, a non-paired test was performed, indicating that the post-hoc models had a better performance for the prediction of all three milk components. Moreover, a similar but paired analysis based exclusively on the samples that were overlapping in the two test sets (*n* = 633) led to the same conclusion.

**Table 4.** Prediction performance statistics for the post-hoc and real-time prediction models

| Model | RMSEP<sup>A</sup> (% w/w) | | | $R_p^2$ | | |
| --- | --- | --- | --- | --- | --- | --- |
|  | Fat | Protein | Lactose | Fat | Protein | Lactose |
| Post-hoc | 0.0784<sup>a</sup> | 0.0797<sup>a</sup> | 0.0772<sup>a</sup> | 0.9892 | 0.9466 | 0.6885 |
| Real-time | 0.0826<sup>b</sup> | 0.1104<sup>b</sup> | 0.0915<sup>b</sup> | 0.9889 | 0.8944 | 0.6442 |

RMSEP = Root-mean-square error of prediction; $R_p^2$ = coefficient of determination.
<sup>A</sup>Within each column, RMSEP-values with different superscripts indicate significant ( $\alpha = 0.05$ ) differences between the different models according to Tukey's HSD multiple comparisons, with a letter lower in the alphabetical order indicating a better model.

The better performance of the post-hoc models compared to the real-time models was hypothesized, as the former models were trained on samples selected over the entire experimental period of 8 weeks, thus better representing the variability present in the samples of the test set. Moreover, fatty acid or amino acid composition of milk can drift over time due to an alteration in the diet or a change in the lactation stage of the cows. Additionally, a shift in the fat globule or casein micelle size or their refractive index can cause changes in the light scattering. If these properties change, they would affect the measured LW-NIR spectra and thus the prediction performance (Aernouts et al., 2015a).

Figure 4 illustrates the drift observed in the prediction errors of (a) fat, (b) protein, and (c) lactose for the test samples by the post-hoc (red) and real-time models (blue). The samples are sorted according to the time when they were measured and they are synchronized for the different models. Because a moving window of 101 points was used to calculate and illustrate the average prediction error and standard deviation over time, with the increase of sample number, these parameters could not be shown for the first and the last 50 samples of the test set. For the post-hoc models, the samples to train the PLSR models are selected over the entire dataset following the duplex approach. Accordingly, the average prediction error and standard deviation of the samples in the test set are interrupted with white zones representing the samples used for calibration. For the real-time models, on the other hand, the calibration samples are all taken at the beginning of the dataset (first week), appearing as a distinct white zone, and thus the test set is a single uninterrupted series of samples.

**Figure 4.**
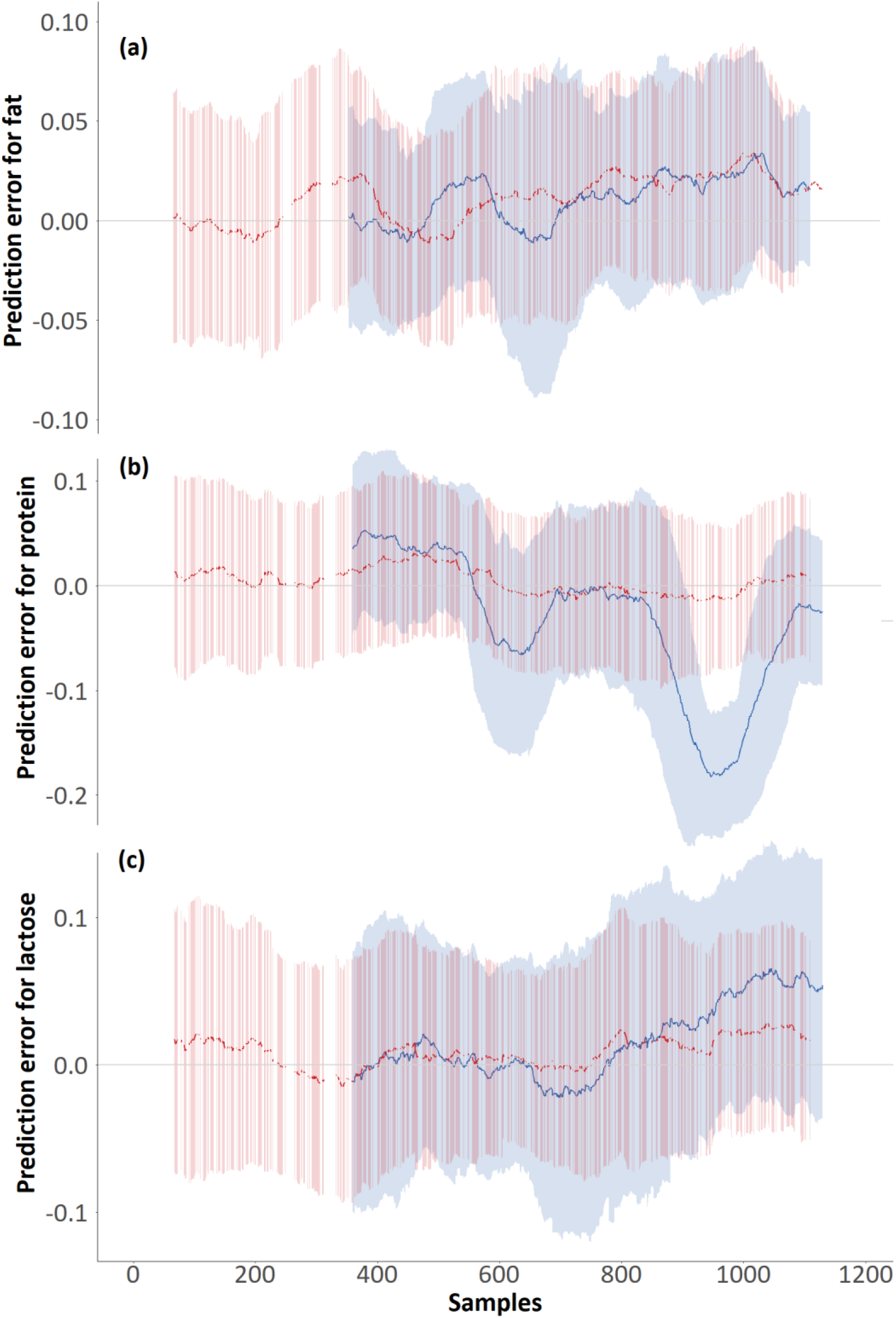
Prediction error plot for (a) fat, (b) protein, and (c) lactose, for the post-hoc (red) and real-time (blue) models, for milk samples of the test set that are ordered based on the time when they were measured. A moving average (101 points) has been applied to obtain the average prediction error (bold line) and the moving standard deviation (red and blue zones, for the post-hoc and real-time models, respectively). The samples selected for calibration are not plotted (white zones).

For protein and lactose, the prediction error of the post-hoc models remains close to zero, with almost no trend with the increasing sample number. In contrast, milk fat prediction for the post-hoc approach and all milk components of the real-time approach display drift over time. The real-time model for the prediction of milk lactose shows a clear drift after week 6 (samples 845 - 1165) with mainly positive prediction errors, indicating an underestimation of the lactose content by the model. The real-time model to predict the milk protein shows apparent drift over the entire test set, although not being very consistent. In weeks 3 and 4 (samples 539 to 677) and 7 (samples 941 – 1108), the prediction errors were mainly negative, indicating an overestimation of the actual milk protein concentration by the models, while they were mainly positive in week 2 (samples 349 – 538). The real-time model for the prediction of milk fat resulted in mainly positive prediction errors in week 3 (sample 539 – 617) and after week 5 (sample 678 – 1165), while the post-hoc counterpart resulted in mainly positive prediction errors in week 2 (samples 349 – 538) and after week 4 (samples 618 – 1165). All real-time models show an increased standard deviation compared to their post-hoc counterparts, and this effect is larger for lactose.

As the post-hoc models include calibration samples throughout the experiment, possible changes over time in milk composition, environmental fluctuations, variations in the morphology and optical properties of fat globules and casein micelles are included in the calibration of the model. This sampling procedure contributes to making this approach more robust against the observed drift. In contrast, the real-time models display a more evident degradation of their prediction capabilities over time. As these models are exclusively trained on the samples collected in the first week, they may not have captured this additional complexity.

Table 4 and Figure 4 indicate that the variability of fat, protein, and lactose over this 8-week experiment is not entirely captured by the calibration dataset of the real-time approach. This difference in milk composition prediction between the two scenarios suggests that especially the real-time models can benefit from calibration maintenance strategies to assure that the composition estimates remain accurate under varying conditions. As such, changes due to management, parity or diet that influence the prediction performance could be included. By adding new samples to the calibration set that represent this fluctuation, the models can be made robust against new measurement conditions, resulting in better overall predictions (Wise and Roginski, 2015). The addition of new samples to the calibration set would require the acquisition of additional reference chemical analyses over time. Currently, individual cow milk samples are already collected every 4 to 6 weeks as a part of milk recording programs. In the context of these programs, the milk composition sensor system would increase the monitoring frequency of fat, protein, and lactose, decreasing the need for milk composition reference analysis and its costs. The calibration maintenance could still benefit from these existing programs by requesting the reference analysis of the most informative samples to contribute to the calibration model.

The accuracy of the real-time prediction approach (Table 4) was evaluated against the ICAR requirements for on-farm in-line and at-line milk analyzers and laboratory milk analyzers. Good predictions were achieved for protein, with an RMSEP value close to 0.11% (all % are *w*/*w*), attaining sufficient accuracy for protein predictions according to ICAR recommendations for at-line (0.2%) and inline (0.25%) milk analyses. Very good results were obtained for fat and lactose, with RMSEP values under 0.1%, accomplishing the more restrictive accuracy limit requested for laboratory milk analyses (0.1%).

### Analysis of the cow-specific bias on the milk predictions by the selected real-time model

The relationship between the median residuals of the calibration data and the prediction residuals of the test data for the real-time prediction model are illustrated in Figure 5 for (a) fat, (b) protein, and (c) lactose for each dairy cow. The concordance correlation coefficient indicates practically no correlation for fat and protein, but a weak relationship for lactose.

**Figure 5.**
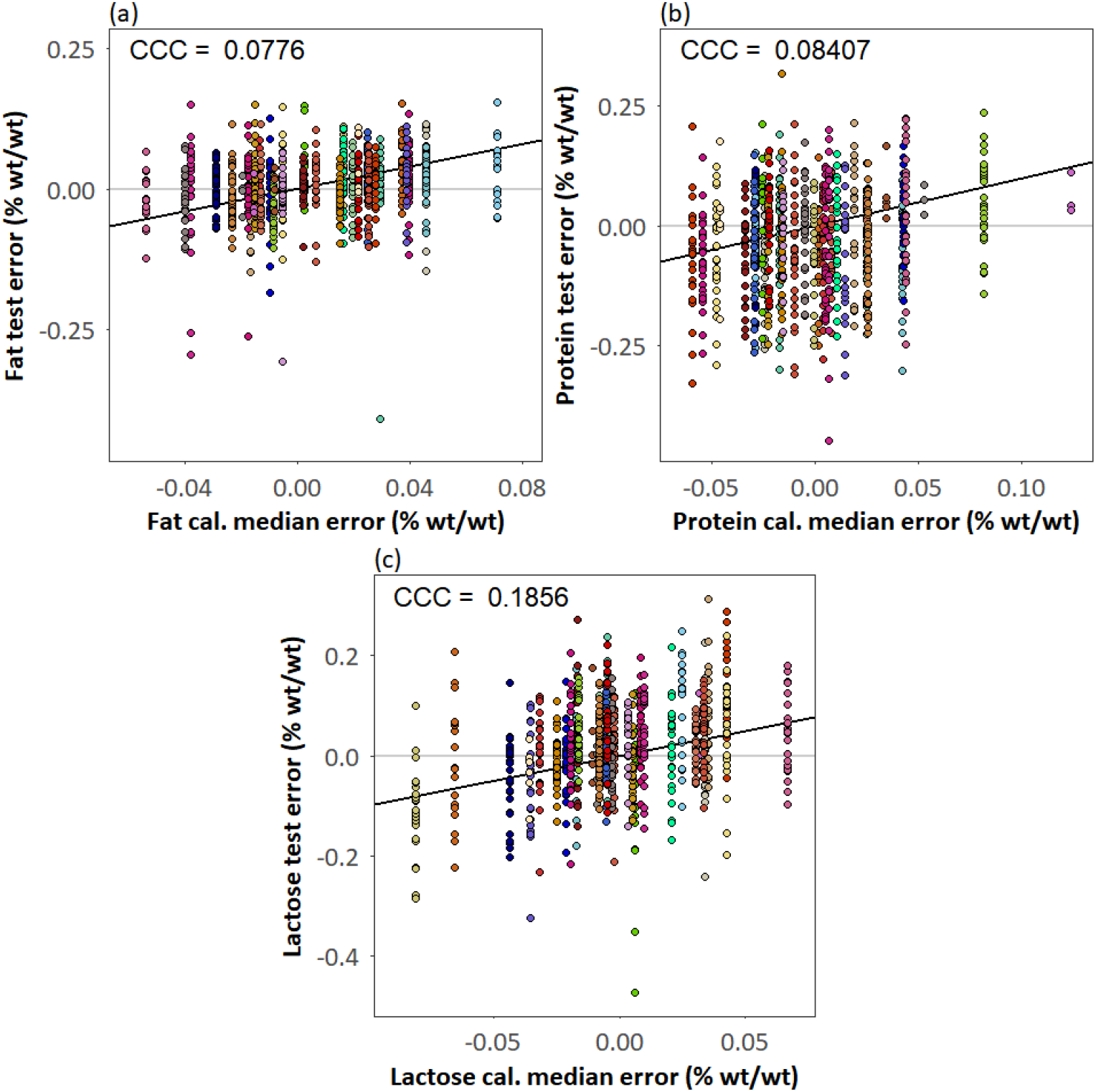
Relationship between prediction residuals and the medians of the calibration residuals for the real-time model, for every individual dairy cow for (a) fat, (b) protein, and (c) lactose. Each color represents the predictions for a single cow. The black solid lines are the identity lines. Three extreme fat test errors were outside the range of figure (a) fat (fat test error of −1.573, −0.661 and −0.556; (% w/w)). CCC = concordance correlation coefficient.

Cow-specific bias correction was performed by subtracting the median residuals of the calibration data for a single cow from all further milk composition predictions in the test set for that same animal. Next, the prediction outcome of the original real-time model and the outcome after applying this cow-specific bias correction were compared with a paired test (Table 6).

**Table 6.** Prediction performance statistics for the real-time and cow-specific bias correction prediction results.

| Model | RMSEP<sup>A</sup> (% w/w) | | | $R_p^2$ | | |
| --- | --- | --- | --- | --- | --- | --- |
|  | Fat | Protein | Lactose | Fat | Protein | Lactose |
| Real-time – No correction | 0.0826 | 0.1104 | 0.0915<sup>b</sup> | 0.9889 | 0.8944 | 0.6442 |
| Real-time – Cow-specific bias correction | 0.0838 | 0.1093 | 0.0875<sup>a</sup> | 0.9886 | 0.8964 | 0.6745 |

RMSEP = Root-mean-square error of prediction; $R_p^2$ = coefficient of determination.
<sup>A</sup>Within each column, RMSEP-values with different superscripts indicate significant ( $\alpha = 0.05$ ) differences between the different models according to Tukey's HSD multiple comparisons, with a letter lower in the alphabetical order indicating a better model.

The cow-specific bias correction resulted in a small improvement in the protein and lactose prediction, being statistically significant for the latter. The RMSEP reduction on lactose increases its performance and brings it closer to the performance of the post-hoc model for this milk component (RMSEP of 0.0772% *w*/*w* and an 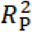 of 0.6885) and further proves its suitability as a laboratory milk composition predictor according to the ICAR guidelines.

Three extreme fat test errors of negative sign (not shown in figure 5), with an absolute value greater than ten times the interquartile range, correspond to three different cows with median errors of positive sign in the calibration set. As the bias correction was performed by subtracting the median residuals of the calibration data from the predictions in the test set, the cow specific correction slightly increased the overall prediction error for fat for all samples. This correction results in a slightly decreased performance compared to not applying any correction for this milk component (Table 6). However, if the sample with the most extreme fat test error (with an absolute value greater than 17 times the interquartile range) is excluded, a very high fat prediction performance (RMSEP of 0.0631% *w*/*w* and a 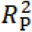 of 0.9935) is obtained for the real-time approach, indicating that an improved overall milk fat prediction can be accomplished. In this new scenario, the cow-specific bias correction for fat further improves (not significantly) the prediction performance, given the newly increased relationship between prediction residuals and the medians of the calibration residuals. However, the NIR spectrum and reference chemical analysis results of this sample were not outlying and no observations made during the trial pointed at valid reasons to justify the removal of this sample. Therefore, this sample was kept in the analysis.

Cow-specific bias correction based on the subtraction of calibration median residuals from the predictions in the test set provided a small, but significant performance increase on the real-time approach for lactose. Therefore, further research is recommended on the application of different correction approaches that more closely reflect the relationship between calibration and test data. Cow-specific bias correction could further improve and be integrated with the aforementioned calibration maintenance procedures.

## Conclusions

An on-farm sensor system for online fat, protein, and lactose analysis was developed and implemented on an AMS. This system can analyze the milk of each individual milking session autonomously to support the cost-efficient monitoring of performance, efficiency, and welfare of individual dairy cows. The prediction accuracy of the system was very high for all three milk components (0.0784% (*w*/*w*), 0.0797% (*w*/*w*) and 0.0772% (*w*/*w*) for fat, protein, and lactose respectively) when creating a prediction model following a post-hoc approach for sample selection. When implemented as a real-time system, training the prediction model with the samples from the first week of the trial and evaluating the model performance on the samples that were measured afterward, the prediction accuracies deteriorated to 0.0826% (*w/w*), 0.1104% (*w/w*) and 0.0915% (*w/w*) for fat, protein, and lactose, respectively. These performances still abide by the ICAR guidelines for on-farm analysis systems for all components and even comply with the ICAR thresholds for laboratory analysis systems for fat and lactose. Additionally, the performance of the real-time model for lactose prediction was further improved with the use of a cow-specific bias correction. As drift was identified in the milk composition predictions over time, future research should focus on the development and implementation of intelligent and efficient calibration maintenance procedures to cope with external influences and guarantee long-term prediction stability.

## Acknowledgments

This work was supported by the government agency Flanders Innovation and Entrepreneurship (VLAIO, Belgium) through projects “Koesensor” (LA110770) and MastiMan (HBC.2016.0774). Milk samples were collected within a trial set-up by ILVO for the “SMART melken” project (LA135081), funded by VLAIO. José A. Diaz Olivares received funding from the Research Foundation Flanders (FWO, Belgium) through grant No. 1S76320N. Ines Adriaens received funding from a KU Leuven (Belgium) postdoctoral mandate grant No. PDM/19/132.

## References

Abd El-Fattah, A.M., Abd Rabo, F.H., EL-Dieb, S.M., Elkashef, H.A.S., 2012. Changes in composition of colostrum of Egyptian buffaloes and Holstein cows. BMC Vet. Res. 8, 19. https://doi.org/10.1186/1746-6148-8-19

Aernouts, B., Adriaens, I., Diaz-Olivares, J., Saeys, W., Mäntysaari, P., Kokkonen, T., Mehtiö, T., Kajava, S., Lidauer, P., Lidauer, M.H., Pastell, M., 2019. Mid-Infrared Spectroscopic Analysis of Raw Milk to Predict the Blood Plasma Non-Esterified Fatty Acid Concentration in Dairy Cows. bioRxiv 853127. https://doi.org/10.1101/853127

Aernouts, B., Polshin, E., Lammertyn, J., Saeys, W., 2011. Visible and near-infrared spectroscopic analysis of raw milk for cow health monitoring: Reflectance or transmittance? J. Dairy Sci. 94, 5315–5329. https://doi.org/10.3168/jds.2011-4354

Aernouts, B., Van Beers, R., Watté, R., Huybrechts, T., Jordens, J., Vermeulen, D., Van Gerven, T., Lammertyn, J., Saeys, W., 2015a. Effect of ultrasonic homogenization on the Vis/NIR bulk optical properties of milk. Colloids Surfaces B Biointerfaces 126, 510–519. https://doi.org/10.1016/j.colsurfb.2015.01.004

Aernouts, B., Van Beers, R., Watté, R., Huybrechts, T., Lammertyn, J., Saeys, W., 2015b. Visible and near-infrared bulk optical properties of raw milk. J. Dairy Sci. 98, 6727–6738. https://doi.org/10.3168/jds.2015-9630

Andersen, H.V., Wedelsback, H., Hansen, P.W., 2013. A white paper from FOSS: NIR spectrometer technology comparison. FOSS P/N 1026672, 1–14.

Anderson, G.P.S., Zhang, I.L., Winkelman, A.M., Harris, B.L., 2017. Comparison of records from in-line milk meters and conventional herd testing for management and genetic evaluation of dairy cows, in: ICAR Technical Series. pp. 203–209.

Bobbo, T., Cipolat-Gotet, C., Bittante, G., Cecchinato, A., 2016. The nonlinear effect of somatic cell count on milk composition, coagulation properties, curd firmness modeling, cheese yield, and curd nutrient recovery. J. Dairy Sci. 99, 5104–5119. https://doi.org/10.3168/jds.2015-10512

Bogomolov, A., Melenteva, A., 2013. Scatter-based quantitative spectroscopic analysis of milk fat and total protein in the region 400-1100nm in the presence of fat globule size variability. Chemom. Intell. Lab. Syst. 126, 129–139. https://doi.org/10.1016/j.chemolab.2013.02.006

Brandt, M., Haeussermann, A., Hartung, E., 2010. Invited review: Technical solutions for analysis of milk constituents and abnormal milk. J. Dairy Sci. https://doi.org/10.3168/jds.2009-2565

Cederkvist, H.R., Aastveit, A.H., Næs, T., 2005. A comparison of methods for testing differences in predictive ability. J. Chemom. 19, 500–509. https://doi.org/10.1002/cem.956

Fadul-Pacheco, L., Lacroix, R., Séguin, M., Grisé, M., Vasseur, E., Lefebvre, D., 2018. Comparison of milk composition and somatic cell count estimates from automatic milking systems sensors and milk recording laboratory, in: Proceedings of the 42nd ICAR Conference. Auckland, New Zealand.

Forsbäck, L., Lindmark-Månsson, H., Andrén, A., Åkerstedt, M., Andrée, L., Svennersten-Sjaunja, K., 2010. Day-to-day variation in milk yield and milk composition at the udder-quarter level. J. Dairy Sci. 93, 3569–3577. https://doi.org/10.3168/jds.2009-3015

Forsbäck, L., Lindmark-Månsson, H., Åndrén, A., Kerstedt, M., Svennersten-Sjaunja, K., 2009. Udder quarter milk composition at different levels of somatic cell count in cow composite milk. Animal 3, 710–717. https://doi.org/10.1017/S1751731109004042

Gonçalves, J.L., Tomazi, T., Barreiro, J.R., Beuron, D.C., Arcari, M.A., Lee, S.H.I., Martins, C.M. de M.R., Araújo Junior, J.P., Santos, M.V. dos, 2016. Effects of bovine subclinical mastitis caused by Corynebacterium spp. on somatic cell count, milk yield and composition by comparing contralateral quarters. Vet. J. 209, 87–92. https://doi.org/10.1016/j.tvjl.2015.08.009

Haaland, D.M., Thomas, E. V., 1988. Partial Least-Squares Methods for Spectral Analyses. 1. Relation to Other Quantitative Calibration Methods and the Extraction of Qualitative Information. Anal. Chem. 60, 1193–1202. https://doi.org/10.1021/ac00162a020

ISO, 2013. Milk and liquid milk products -- Guidelines for the application of mid-infrared spectrometry. Page 14 in International Standard ISO 9622:2013/IDF 141:2013. International Dairy Federation.

Kaniyamattam, K., De Vries, A., 2014. Agreement between milk fat, protein, and lactose observations collected from the Dairy Herd Improvement Association (DHIA) and a real-time milk analyzer. J. Dairy Sci. 97, 2896–2908. https://doi.org/10.3168/jds.2013-7690

Kawamura, S., Kawasaki, M., Nakatsuji, H., Natsuga, M., 2007. Near-infrared spectroscopic sensing system for online monitoring of milk quality during milking. Sens. Instrum. Food Qual. Saf. 1, 37–43. https://doi.org/10.1007/s11694-006-9001-x

Kawasaki, M., Kawamura, S., Tsukahara, M., Morita, S., Komiya, M., Natsuga, M., 2008. Near-infrared spectroscopic sensing system for on-line milk quality assessment in a milking robot. Comput. Electron. Agric. 63, 22–27. https://doi.org/10.1016/j.compag.2008.01.006

Kemps, B.J., Saeys, W., Mertens, K., Darius, P., De Baerdemaeker, J.G., De Ketelaere, B., 2010. The importance of choosing the right validation strategy in inverse modelling. J. Near Infrared Spectrosc. 18, 231–237. https://doi.org/10.1255/jnirs.882

Lin, L.I.-K., 1989. A Concordance Correlation Coefficient to Evaluate Reproducibility. Biometrics 45, 255. https://doi.org/10.2307/2532051

Mäntysaari, P., Mäntysaari, E.A., Kokkonen, T., Mehtiö, T., Kajava, S., Grelet, C., Lidauer, P., Lidauer, M.H., 2019. Body and milk traits as indicators of dairy cow energy status in early lactation. J. Dairy Sci. 102, 7904–7916. https://doi.org/10.3168/jds.2018-15792

McParland, S., Lewis, E., Kennedy, E., Moore, S.G., McCarthy, B., O’Donovan, M., Butler, S.T., Pryce, J.E., Berry, D.P., 2014. Mid-infrared spectrometry of milk as a predictor of energy intake and efficiency in lactating dairy cows. J. Dairy Sci. 97, 5863–5871. https://doi.org/10.3168/jds.2014-8214

Melfsen, A., Hartung, E., Haeussermann, A., 2012. Accuracy of in-line milk composition analysis with diffuse reflectance near-infrared spectroscopy. J. Dairy Sci. 95, 6465–6476. https://doi.org/10.3168/jds.2012-5388

R Core Team, 2018. R: A language and environment for statistical computing. R Foundation for Statistical Computing, Vienna, Austria. URL https://www.R-project.org/.

Rosati, A., Van Vleck, L.D., 2002. Estimation of genetic parameters for milk, fat, protein and mozzarella cheese production for the Italian river buffalo Bubalus bubalis population. Livest. Prod. Sci. 74, 185–190. https://doi.org/10.1016/S0301-6226(01)00293-7

Saeys, W., Beullens, K., Lammertyn, J., Ramon, H., Naes, T., 2008. Increasing robustness against changes in the interferent structure by incorporating prior information in the augmented classical least-squares framework. Anal. Chem. 80, 4951–4959. https://doi.org/10.1021/ac800155n

Saranwong, S., Kawano, S., 2008. System design for non-destructive near infrared analyses of chemical components and total aerobic bacteria count of raw milk. J. Near Infrared Spectrosc. 16, 389–398. https://doi.org/10.1255/jnirs.807

Shetty, N., Min, T.G., Gislum, R., Olesen, M.H., Boelt, B., 2011. Optimal sample size for predicting viability of cabbage and radish seeds based on near infrared spectra of single seeds. J. Near Infrared Spectrosc. 19, 451–461. https://doi.org/10.1255/jnirs.966

Slutsky, B., 1997. Handbook of Chemometrics and Qualimetrics: Part A, Data Handling in Science and Technology Volume 20A. Elsevier, Amsterdam. https://doi.org/10.1021/ci980427d

Snee, R.D., 1977. Validation of Regression Models: Methods and Examples. Technometrics 19, 415–428. https://doi.org/10.1080/00401706.1977.10489581

Tsenkova, R., Atanassova, S., Ozaki, Y., Toyoda, K., Itoh, K., 2001. Near-infrared spectroscopy for biomonitoring: Influence of somatic cell count on cow’s milk composition analysis. Int. Dairy J. 11, 779–783. https://doi.org/10.1016/S0958-6946(01)00110-8

Tsenkova, R., Atanassova, S., Toyoda, K., Ozaki, Y., Itoh, K., Fearn, T., 1999. Near-infrared spectroscopy for dairy management: Measurement of unhomogenized milk composition. J. Dairy Sci. 82, 2344–2351. https://doi.org/10.3168/jds.S0022-0302(99)75484-6

Wise, B.M., Roginski, R.T., 2015. A calibration model maintenance roadmap, in: IFAC-PapersOnLine. pp. 260–265. https://doi.org/10.1016/j.ifacol.2015.08.191

Zamora-Rojas, E., Aernouts, B., Garrido-Varo, A., Pérez-Marín, D., Guerrero-Ginel, J.E., Saeys, W., 2013. Double integrating sphere measurements for estimating optical properties of pig subcutaneous adipose tissue. Innov. Food Sci. Emerg. Technol. 19, 218–226. https://doi.org/10.1016/j.ifset.2013.04.015

