## Supplementary Materials (Tables S1-S4) for "Online milk composition analysis with an on-farm near-infrared sensor"

To improve milk composition predictions and decrease degradation caused by spectral drift, different time intervals for taking into account a new pair of dark and white reference spectra were evaluated. Five different transmittance datasets were obtained by considering time intervals of 0.5, 1 and 3 hours between two spectral reference pairs, by only regarding a new pair of spectral references at the start of each session after connecting (AC) the sensor system to the AMS and by taking a new dark and white spectrum for each sample measurement (time interval of 0 hours). For each of the five transmittance data sets, a post-hoc and real-time prediction model for each milk component was trained and tested. The procedures used for obtaining an optimal reference acquisition interval are detailed in this section.

#### *Selection of the best time interval for dark and white spectral references for the post-hoc approach*

For each of the three milk components, five post-hoc models were obtained, each representing a different time interval for taking a new pair of dark and white spectral references. The optimal time interval was selected for each milk component following the same procedure, as described in the *Materials and Methods* section, to select the number of latent variables for a PLSR model, the best spectral preprocessing, and the most informative spectral wavelengths. Moreover, we selected the time interval that resulted in the least complex PLSR model (with the lowest number of latent variables) for which the cross-validation predictions were not significantly worse compared to the model with the lowest root-mean-square error of cross-validation (RMSECV). These comparisons were based on one-sided paired *T*-tests ($\alpha$ = 0.05) applied to the absolute residuals of the cross-validated calibration samples (Cederkvist et al., 2005).

For each milk component, the five models were applied to the test subset of the five respective transmittance datasets. Accordingly, milk composition predictions were obtained for the samples in the test set, and the respective residuals were calculated. These errors were then combined into a root-mean-square error of prediction (RMSEP) for each of the five models. Finally, the prediction performances of the five models were compared using a two-way repeated-measures analysis of variance (ANOVA) performed on the squared residuals of the samples in the test set. For this procedure, the time interval for taking a new pair of dark and white spectral references was treated as a fixed effect with five levels, while the sample number was specified as a random effect (Cederkvist et al., 2005). Only when a significant difference between the models was detected by the ANOVA procedure ($\alpha$ = 0.05), then the performance of the models was compared mutually with a Tukey's honestly significant difference (HSD) multiple comparison test ($\alpha$ = 0.05).

The number of selected latent variables for each model of each milk component and the respective cross-validation performances in terms of RMSECV and $R^2_{\mathrm{CV}}$ are provided in Table S1. For fat, no significant difference was found between the five post-hoc models based on the cross-validation residuals. Moreover, all models had three latent variables except for the one where a new pair of dark and white spectral references was taken every three hours, which selected six latent variables. Taking a new pair of spectral references every hour resulted in the worst protein prediction, while all other models were not significantly different from each other. From these models, the one for which a new pair of spectral references were taken every half an hour had the lowest number of latent variables. This frequency of taking a new spectral reference pair also resulted in the best lactose predictions, which was significantly better than the other four models. Following the selection procedure, this frequency (AC + every 0.5 hours) and the respective model was selected as the optimal post-hoc model.

It is expected that if new spectral references are taken with a higher frequency than this optimum, unnecessary stochastic noise, typical of a spectral measurement, would be introduced. In contrast, if new spectral references are taken with a lower frequency than this optimum, significant drift in the spectral output of the light source and spectral sensitivity of the spectrometers more influential than this stochastic noise might not be fully captured and accounted for, also negatively affecting the results.

**Table S1.** Statistics indicating the performance of the five post-hoc models, representing different time intervals for taking a new pair of dark and white spectral references based on cross-validation of the samples in the calibration set. Additionally, also the complexity of the partial least squares regression models in terms of the number of latent variables is presented.

| New reference | Latent variables | | | RMSECV[Δ] (% *w/w*) | | | $R^2_{CV}$ | | |
|---|---|---|---|---|---|---|---|---|---|
| | Fat | Protein | Lactose | Fat | Protein | Lactose | Fat | Protein | Lactose |
| Every sample | 3 | 12 | 9 | 0.0847 | 0.064[a] | 0.0882[b] | 0.9916 | 0.9637 | 0.7742 |
| AC + every 0.5 hours | 3 | 10 | 16 | 0.0852 | 0.0683[a] | 0.0777[a] | 0.9915 | 0.9588 | 0.8247 |
| AC + every hour | 3 | 8 | 2 | 0.0864 | 0.0726[b] | 0.0848[b] | 0.9912 | 0.9534 | 0.7911 |
| AC + every 3 hours | 6 | 17 | 7 | 0.0858 | 0.0602[a] | 0.0916[b] | 0.9914 | 0.9679 | 0.7563 |
| Only AC | 3 | 19 | 19 | 0.0866 | 0.0575[a] | 0.0910[b] | 0.9912 | 0.9708 | 0.7598 |

AC = after connection of the system to the AMS; RMSECV = Root-mean-square error of cross-validation;

$R^2_{CV}$ = coefficient of determination.

[Δ]Within each column, RMSECV-values with different superscripts indicate significant ($\alpha$ = 0.05) differences between the different models according to Tukey's HSD multiple comparisons, with a letter lower in the alphabetical order indicating a better model.

The selected post-hoc model, corresponding to taking a new spectral reference pair every half hour, resulted in the best lactose predictions for the samples in the test set (Table S2). The predictions of the fat and protein concentration for these samples by the selected post-hoc model were not the best. However, they were not significantly worse compared to the best performing post-hoc models for these respective milk components either.

**Table S2.** Statistics indicating the performance of the five post-hoc models, representing different time intervals for taking a new pair of dark and white spectral references, based on the samples in the test set.

| New reference | RMSEP$^{\Delta}$ (% w/w) | | | $R^2_P$ | | |
|---|---|---|---|---|---|---|
| | Fat | Protein | Lactose | Fat | Protein | Lactose |
| Every sample | 0.0790$^{b}$ | 0.0760$^{a,b}$ | 0.0812$^{a}$ | 0.9892 | 0.9515 | 0.6553 |
| AC + every 0.5 hours | 0.0784$^{a,b}$ | 0.0797$^{a,b}$ | 0.0772$^{a}$ | 0.9892 | 0.9466 | 0.6885 |
| AC + every hour | 0.0808$^{b}$ | 0.0832$^{b}$ | 0.0897$^{b}$ | 0.9886 | 0.9419 | 0.5789 |
| AC + every 3 hours | 0.0771$^{a}$ | 0.0796$^{a,b}$ | 0.0897$^{b}$ | 0.9896 | 0.9468 | 0.5796 |
| Only AC | 0.0783$^{a,b}$ | 0.0714$^{a}$ | 0.0783$^{a}$ | 0.9893 | 0.9572 | 0.6794 |

AC = after connection of the system to the AMS; RMSEP = Root-mean-square error of prediction;

$R^2_P$ = coefficient of determination.

$^{\Delta}$Within each column, RMSEP-values with different superscripts indicate significant ($\alpha$ = 0.05) differences between the different models according to Tukey's HSD multiple comparisons, with a letter lower in the alphabetical order indicating a better model.

### *Selection of the best time interval for dark and white spectral references for the real-time approach*

For each milk component, Table S3 reports the complexity (i.e. number of latent variables of the PLSR models) and the performance statistics for the five real-time models, optimized for the cross-validation of the samples in the calibration set. Similar to the post-hoc models, no significant difference was found between the five real-time models when evaluating the prediction of the fat content. The least complex fat prediction models were the ones that considered a new spectral reference pair every half hour and every hour, having selected only four latent variables. Also, the protein prediction models were performing not significantly different, except when a new spectral reference pair was taken for every sample measurement, which resulted in significantly worse predictions. From the remaining four models, the one for which a new spectral reference pair was taken every half an hour was the least complex with only five selected latent variables. Also, for the prediction of lactose, this frequency of taking a new spectral reference pair resulted in the least complex model that did not perform significantly worse compared to the real-time model with the lowest RMSECV.

From this analysis, it was concluded that taking a new spectral reference pair every half an hour (AC + every 0.5 hours) was the best, and the respective model for each milk component was thus selected as the optimal real-time model. Although only a few significant differences were found between the five real-time models, this optimal frequency of taking a new spectral reference pair resulted in the least complex prediction models. It is hypothesized that all other frequencies of reference acquisition result in additional latent variables taken up by the PLSR models to account for the additional noise or drift that is being introduced.

**Table S3.** Prediction performance and model fit of the five real-time models, representing different time intervals for taking a new pair of dark and white spectral references based on cross-validation of the samples in the calibration set. Additionally, also the complexity of the partial least squares regression models in terms of the number of latent variables is presented.

| New reference | Latent variables | | | RMSECV[Δ] (% w/w) | | | $R^2_{CV}$ | | |
|---|---|---|---|---|---|---|---|---|---|
| | Fat | Protein | Lactose | Fat | Protein | Lactose | Fat | Protein | Lactose |
| Every sample | 9 | 7 | 20 | 0.0614 | 0.0747[b] | 0.0610 | 0.9949 | 0.9546 | 0.8412 |
| AC + every 0.5 hour | 4 | 5 | 13 | 0.0660 | 0.0668[a] | 0.0687 | 0.9941 | 0.9636 | 0.7989 |
| AC + every hour | 4 | 20 | 16 | 0.0662 | 0.0634[a] | 0.0646 | 0.9941 | 0.9673 | 0.8219 |
| AC + every 3 hours | 7 | 19 | 14 | 0.0648 | 0.0591[a] | 0.0678 | 0.9943 | 0.9715 | 0.8038 |
| Only AC | 18 | 15 | 19 | 0.0624 | 0.0525[a] | 0.0666 | 0.9947 | 0.9775 | 0.8109 |

AC = after connection of the system to the AMS; RMSECV = Root-mean-square error of cross-validation;

$R^2_{CV}$ = coefficient of determination.

[Δ]Within each column, RMSECV-values with different superscripts indicate significant ($\alpha$ = 0.05) differences between the different models according to Tukey's HSD multiple comparisons, with a letter lower in the alphabetical order indicating a better model.

The selected real-time model for each milk component, corresponding to taking a new spectral reference pair every half an hour, resulted in the best fat predictions for the samples in the test set (Table S4). The predictions of the protein and lactose concentration for these samples by the selected real-time model were not the best. However, they were not significantly worse compared to the best performing real-time models for these respective milk components either.

**Table S4.** Statistics indicating the performance of the five real-time models, representing different time intervals for taking a new pair of dark and white spectral references, based on the samples in the test set.

| New reference | RMSEP[Δ] (% w/w) | | | $R^2_P$ | | |
|---|---|---|---|---|---|---|
| | Fat | Protein | Lactose | Fat | Protein | Lactose |
| Every sample | 0.1006[b] | 0.1085[a] | 0.1207[b] | 0.9836 | 0.8980 | 0.3807 |
| AC + every 0.5 hours | 0.0826[a] | 0.1104[a] | 0.0915[a] | 0.9889 | 0.8944 | 0.6442 |
| AC + every hour | 0.0843[a] | 0.1186[a,b] | 0.1147[b] | 0.9885 | 0.8781 | 0.4405 |
| AC + every 3 hours | 0.0868[a] | 0.1215[b] | 0.0897[b] | 0.9878 | 0.8722 | 0.6576 |
| Only AC | 0.0983[b] | 0.1080[a] | 0.1936[a] | 0.9843 | 0.8989 | 0 |

AC = after connection of the system to the AMS; RMSEP = Root-mean-square error of prediction;

$R^2_P$ = coefficient of determination.

[Δ]Within each column, RMSEP-values with different superscripts indicate significant ($\alpha$ = 0.05) differences between the different models according to Tukey's HSD multiple comparisons, with a letter lower in the alphabetical order indicating a better model.
